# BIMSA: Accelerating Long Sequence Alignment Using Processing-In-Memory

**DOI:** 10.1101/2024.05.10.593513

**Authors:** Alejandro Alonso-Marín, Ivan Fernandez, Quim Aguado-Puig, Juan Gómez-Luna, Santiago Marco-Sola, Onur Mutlu, Miquel Moreto

## Abstract

**Motivation:** Recent advances in sequencing technologies have stressed the critical role of sequence analysis algorithms and tools in genomics and healthcare research. In particular, sequence alignment is a fundamental building block in many sequence analysis pipelines and is frequently a performance bottleneck both in terms of execution time and memory usage. Classical sequence alignment algorithms are based on dynamic programming and often require quadratic time and memory with respect to the sequence length. As a result, classical sequence alignment algorithms fail to scale with increasing sequence lengths and quickly become memory-bound due to data-movement penalties.

**Results:** Processing-In-Memory (PIM) is an emerging architectural paradigm that seeks to accelerate memory-bound algorithms by bringing computation closer to the data to mitigate data-movement penalties. This work presents BIMSA (<u>B</u>idirectional In-<u>M</u>emory <u>S</u>equence <u>A</u>lignment), a PIM design and implementation for the state-of-the-art sequence alignment algorithm BiWFA (Bidirectional Wavefront Alignment), incorporating new hardware-aware optimizations for a production-ready PIM architecture (UPMEM). BIMSA supports aligning sequences up to 100K bases, exceeding the limitations of state-of-the-art PIM implementations. First, BIMSA achieves speedups up to 22.24*×* (11.95*×* on average) compared to state-of-the-art PIM-enabled implementations of sequence alignment algorithms. Second, achieves speedups up to 5.84*×* (2.83*×* on average) compared to the highest-performance multicore CPU implementation of BiWFA. Third, BIMSA exhibits linear scalability with the number of compute units in memory, enabling further performance improvements with upcoming PIM architectures equipped with more compute units and achieving speedups up to 9.56*×* (4.7*×* on average).

**Availability:** Code and documentation are publicly available at https://github.com/AlejandroAMarin/BIMSA.

## 1 Introduction

The alignment of DNA, RNA, or protein sequences is essential for understanding various biological processes and identifying genetic variations (Alser *et al*., 2022; Churko *et al*., 2013; Roy *et al*., 2018). Due to advances in sequencing technologies, sequences produced are becoming longer, reaching more than 1 M bases in some cases (Reuter *et al*., 2015; Schloss, 2008).

Sequence alignment based on Dynamic Programming (DP), such as Needleman-Wunsch (Needleman and Wunsch, 1970) or Smith-Waterman (Waterman *et al*., 1976; Gotoh, 1982) algorithms, is widely used and effective for aligning short sequences. However, it presents certain challenges, especially when dealing with ultra-long sequences or large-scale data sets. In particular, classical DP-based algorithms present a time and space complexity that grows quadratically *O*(*n*^2^) (where *n* is the length of the sequences), requiring both computation and storage of large matrices to track partial solutions during the alignment process (Alser *et al*., 2021). As a result, these algorithms cannot scale when aligning long-sequence datasets generated by modern sequencing technologies (Marco-Sola *et al*., 2021).

To address these challenges, (Marco-Sola *et al*., 2021) introduced WFA, a novel algorithm that takes advantage of homologous regions between sequences to accelerate alignment computation. In essence, WFA computes optimal alignments in *O*(*ns*) time and *O*(*s*^2^) memory, where *n* is the sequence length and *s* is the optimal alignment score. While WFA reduces computational complexity and memory footprint, prior work (Diab *et al*., 2023) has demonstrated that sequence alignment algorithms, including the WFA algorithm, are memory-bound due to their low arithmetic intensity and large memory footprint. As a result, existing hardware acceleration solutions based on CPUs (Balhaf *et al*., 2016; Daily, 2016; Vasimuddin *et al*., 2019; Hajinazar *et al*., 2021), GPUs (Aguado-Puig *et al*., 2023, 2022; Gerometta *et al*., 2023), FPGAs (Haghi *et al*., 2021b,a) and ASICs (Haghi *et al*., 2023; Walia *et al*., 2024)) spend most of their execution time waiting for memory requests to be served, frequently leaving their powerful compute units idle.

Recently, (Marco-Sola *et al*., 2023) introduced the BiWFA algorithm, which improves on the original WFA algorithm by reducing memory space to *O*(*s*) and maintaining compute time at *O*(*ns*). Nevertheless, our performance characterization of BiWFA reveals that it remains a memory-bound algorithm whose performance is bottlenecked by data movement between memory and compute units, especially when aligning long and noisy sequences.

To overcome such memory bottlenecks, Processing-In-Memory (PIM) (Kautz, 1969; Stone, 1970; Mutlu *et al*., 2019a,b; Ghose *et al*., 2019; Mutlu *et al*., 2022) has emerged as a novel computing paradigm designed to alleviate data-movement penalties by placing compute capabilities closer to where data is stored. PIM devices are increasingly becoming available in the commercial market. A notable example is the UPMEM PIM architecture (UPMEM, 2024; Gómez-Luna *et al*., 2022; Gómez-Luna *et al*., 2021), which is based on DDR4 DRAM memory. The UPMEM architecture places small general-purpose processors, called *DPUs* (DRAM Processing Units), near the DRAM banks inside each DRAM chip. As a result, the DPUs access memory at lower latency and higher bandwidth than conventional processors (e.g., CPU). The recently presented Alignment-In-Memory (AIM) (Diab *et al*., 2023) library explores the acceleration of WFA on UPMEM PIM devices. Despite its performance improvement, our analysis reveals that AIM’s performance is limited by the *O*(*s*^2^) memory space requirements when aligning sufficiently long and noisy sequences (i.e., highly dissimilar sequences containing errors and gaps).

Our **goal** in this work is to *enable fast alignment of sequences of different lengths*, ranging from short sequences to ultra-long sequences (e.g., 100 K bps) exploiting PIM. To this end, we present BIMSA (Bidirectional In-Memory Sequence Alignment), the first PIM-enabled memory-efficient alignment design and implementation which exploits the linear memory complexity of BiWFA to efficiently perform sequence alignment in the DPUs of the UPMEM PIM system. In summary, this work makes the following contributions.

- We characterize CPU-based state-of-the-art sequence alignment algorithms, including WFA (Marco-Sola *et al*., 2021) and BiWFA (Marco-Sola *et al*., 2023), to demonstrate that they are bottlenecked by memory. Moreover, we thoroughly characterize the Alignment-In-Memory (AIM) library implementation of WFA (Diab *et al*., 2023) and identify its limitations for aligning long sequences.
- We design and implement a PIM-enabled version of BiWFA tailored for the UPMEM architecture, incorporating PIM-aware optimizations to maximize performance. We show that our new design can compute optimal alignments of sequence lengths longer than what is possible with AIM, the state-of-the-art PIM design.
- We evaluate BIMSA’s performance for different parameter configurations and compare it with the AIM library and BiWFA’s CPU implementation in the WFA2-lib (Marco-Sola *et al*., 2023). Compared to the AIM library, BIMSA provides performance speedups up to 22.24*×* (11.95*×* on average). When compared to CPU WFA2-lib, our solution increases performance up to 5.84*×* (2.83*×* on average). Additionally, we show a performance projection for the upcoming UPMEM-v1B system chips, achieving speedups up to 9.56*×* (4.7*×* on average) compared to WFA2-lib (Marco-Sola *et al*., 2023).
- We discuss PIM’s strengths and limitations for sequence alignment when processing real sequencing datasets and how future PIM systems can overcome such limitations in future generations of the technology.

## 2 Background and Motivation

### 2.1 Sequence Alignment Algorithms

Let *Q* and *R* be two sequences (query *Q* and reference *R*) of length *n* and *m* respectively, and *{M, X, I, D}* a set of penalty scores, corresponding to the cost of match, mismatch, insertion, and deletion. Classical Dynamic Programming (DP) algorithms, like Needleman-Wunsch (Needleman and Wunsch, 1970), calculate the score of aligning all suffixes between two sequences stored in an *n × m* matrix using simple operations. Calculating the DP-matrix results in a *O*(*nm*) complexity in compute and space. The operations that represent the calculation of a single DP-matrix cell *M*_*i,j*_ are shown in Eq. 1.

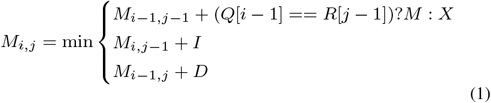

The WFA, introduced by (Marco-Sola *et al*., 2021), is a sequence alignment algorithm that computes cells of the DP-matrix in increasing order of score. It utilizes a data structure *W*, called *Wavefront*, which keeps track of the most advanced cell on each diagonal for a certain score. Wavefronts dynamically expand as the alignment progress continues until its completion. Their computation involves two main steps, known as *Compute Wavefront* and *Extend Wavefront*:

#### (1) Compute Wavefront

This step determines each new value of the next wavefront by evaluating the current minimum value among adjacent cells. Each cell calculation follows Eq. 2.

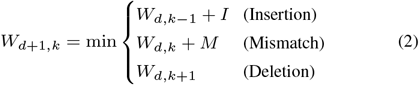

#### (2) Extend Wavefront

This step comprises the computation of the Longest Common Prefix (LCP) for each diagonal of the wavefront. A single cell extend is represented in Eq. 3.

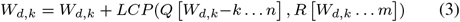

WFA improves classic DP-based algorithms’ complexity from *O*(*n*^2^) to *O*(*ns*) in time, and from *O*(*n*^2^) to *O*(*s*^2^) in memory (where *s* represents the optimal alignment score). Nonetheless, as demonstrated in (Diab *et al*., 2023), WFA suffers from excessive data movement between the memory and the compute units, making it a memory-bound algorithm, similar to traditional DP-based algorithms.

BiWFA (Marco-Sola *et al*., 2023) represents an improvement over the original WFA designed to address memory limitations by reducing memory complexity from *O*(*s*^2^) to *O*(*s*) while retaining the same time complexity *O*(*ns*). The key idea behind BiWFA is to simultaneously align the sequences forward and backward using WFA until the two alignments overlap in the middle. This convergence point is referred to as the *breakpoint*, which is necessarily a point belonging to the optimal alignment path. Moreover, this initial breakpoint already provides the optimal alignment score between the two sequences.

### 2.2 BiWFA Performance Limitations

WFA2lib (Marco-Sola *et al*., 2021) is the state-of-the-art BiWFA implementation for CPU. To identify its performance bottlenecks on server-class CPUs, we run experiments aligning four representative datasets obtained from NIST’s Genome in a Bottle (GIAB) (NIST, 2023) as in (Aguado-Puig *et al*., 2023). We perform this characterization on a dual-socket server equipped with two Intel Xeon Silver 4215 CPUs (Intel, 2019) running at 2.50 GHz and 220 GiB of DRAM memory. Each CPU socket has 8 cores and 2 threads per core (totaling 16 hardware threads). Each core is equipped with an L1d and L1i cache (32 KB each), an L2 cache (1 MB), and a shared L3 cache (11 MB).

Figure 1 shows the scalability of BiWFA using different numbers of threads and real datasets containing sequences of different lengths and error rates. The gray dashed line shows the ideal (linear) speedup relative to the sequential version. We observe that WFA2lib’s BiWFA implementation shows poor scaling across all datasets. We consider `PacBio.CSS` as a representative example of this limitation since using 16 threads merely provides a ≈ 4*×* speedup compared to the single-thread execution.

**Fig. 1:**
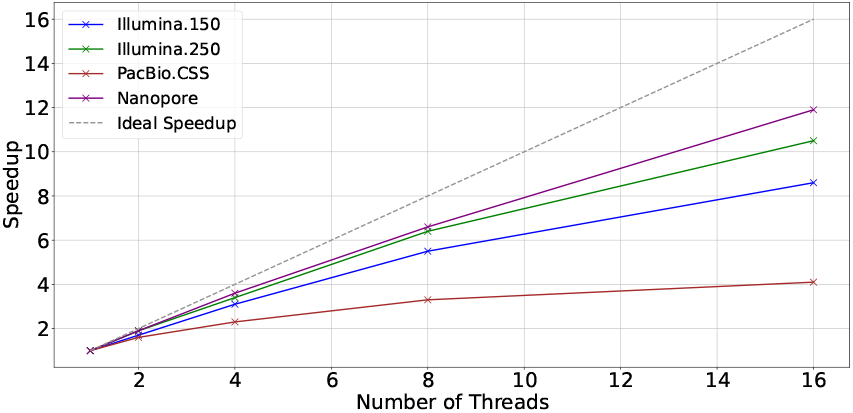
Scalability of WFA2lib’s BiWFA when using a different number of threads on a server with two Intel Xeon Silver 4215 CPUs.

We investigate the cause of BiWFA’s poor scaling on CPUs. Table 1 shows results regarding the percentage of execution cycles CPU spent stalled while the memory subsystem has an outstanding load (i.e., *memory stalls fraction*), the percentage of LLC (Last-Level-Cache) misses, the average load miss latency (i.e., the average number of cycles to serve each load that causes a miss), and the MLP (i.e., a measure of the Memory-Level-Parallelism as the average number of outstanding L1 loads when there is cache miss). For reference, on modern processors, the average latency for accessing L1, L2, LLC, and main memory is 1 cycle, 10 cycles, 50 cycles, and over 200 cycles, respectively. Similarly, MLP values can increase up to 10. Looking at the results in Table 1, we observe that BiWFA executions show high LLC Miss Ratios (> 70%), high-latency loads (7.9 - 56.4 cycles on average), and a high MLP across all experiments, causing BiWFA to be bottlenecked by data movement penalties. In Supplementary Section S2, we include a more detailed performance analysis to reinforce the memory-bound characterization and discuss how the wavefront size and number of working threads affect the CPU execution performance.

**Table 1.** Percentage of Memory Stalls, percentage of LLC Miss, Load Miss Latency, and MLP (Memory Level Parallelism) running WFA2lib’s BiWFA to align real datasets on 16-thread executions.

| Dataset | Memory Stalls (%) | LLC Miss (%) | Load Miss Latency (cycles) | MLP |
| --- | --- | --- | --- | --- |
| Illumina.150 | 21.9 | 95.26 | 56.4 | 7.2 |
| Illumina.250 | 14.7 | 73.22 | 75.0 | 9.2 |
| PacBio.CSS | 21.9 | 86.81 | 38.0 | 9.8 |
| Nanopore | 4.4 | 85.73 | 7.9 | 3.6 |

**Key Observation:** BiWFA presents a high ratio of costly last-level cache misses that prevent scalability. As a consequence, BiWFA is memory-bound on conventional computing platforms.

### 2.3 Processing-In-Memory Paradigm

Common *processor-centric* architectures are equipped with caches that aim to alleviate the gap between processor and memory speeds. Cache hierarchies can sometimes hide long-latency memory accesses when the number of operations per byte (i.e., arithmetic intensity) and locality is high. However, when the application’s arithmetic intensity is low, or there is poor locality, data movement between CPU and memory becomes a bottleneck.

To mitigate this problem, the Processing-In-Memory (PIM) (Mutlu *et al*., 2022) paradigm proposes placing compute units as close to where data is stored, offering higher bandwidth, lower latency, and lower energy consumption than conventional platforms. Although PIM original ideas were introduced decades ago (Kautz, 1969; Stone, 1970), commercial PIM platforms have recently become feasible from a technology standpoint. Notable examples are UPMEM (Gómez-Luna *et al*., 2022; Gómez-Luna *et al*., 2021), Samsung HBM-PIM (Kwon *et al*., 2021; Lee *et al*., 2021), Samsung AxDIMM (Ke *et al*., 2021), SK Hynix AiM (Lee *et al*., 2022), and Alibaba HB-PNM (niu, 2022).

### 2.4 UPMEM PIM Platform Architecture

UPMEM is a commercially available general-purpose PIM platform. Figure 2 shows a high-level view of the UPMEM setup, showing the different memories and how the pipeline is structured. This platform is composed of x86-based host CPUs, *regular* DRAM main memory DIMMs, and PIM-enabled DRAM DIMMs.

**Fig. 2:**
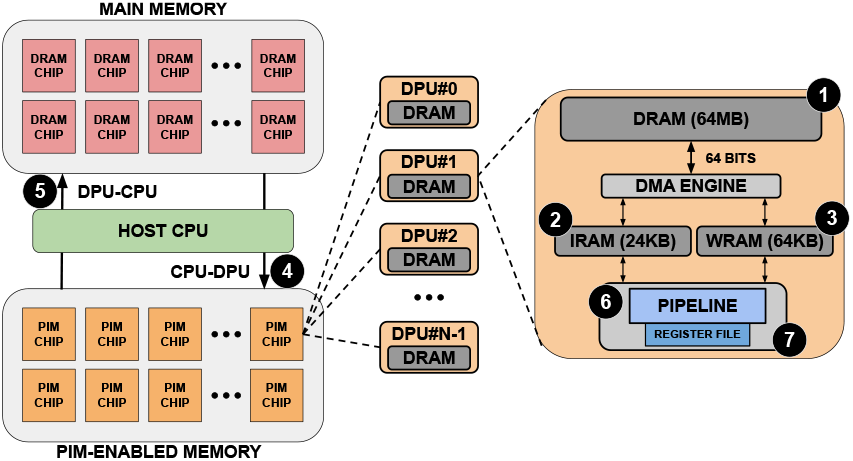
Typical organization of an UPMEM-based server including one or several host CPU(s), main memory, and PIM-enabled memory (left). The internal components of a PIM chip are depicted on the right based on (Gómez-Luna *et al*., 2022).

Each PIM-enabled Memory DIMM is composed of several PIM Chips. Each PIM Chip comprises DRAM Processing Units (DPUs), and each of them is a small general-purpose 32-bit RISC processor that provides 24 hardware threads. Consequently, UPMEM provides two degrees of parallelism: DPUs and threads. Each DPU includes a DRAM bank of 64 MB (MRAM) ❶, 24 KB of Instruction RAM (IRAM) ❷, 64 KB of Working RAM (WRAM) ❸, a register file ❻ and a compute pipeline ❼. Each PIM Chip can be accessed from the host via the DRAM interface, both for data transfer and compute/control via the Control/Status Interface. When data is transferred to PIM-enabled Memory, it is stored in MRAM. DPUs cannot operate directly on the MRAM data, so the DMA Engine moves data to the WRAM, and the instructions to operate over that data are sent to the IRAM.

In a typical PIM execution flow, the initial step involves reading a block of data from MRAM, with the block size specified by the programmer. Subsequently, this block is allocated within a pre-defined WRAM data structure. Following the WRAM data allocation, the general-purpose processor gains access to the WRAM data as an array and can perform computations as needed. Upon completion of the computation, the processed data is written back to MRAM in the same block size. It is important to note that all data transfers to MRAM must be 8-byte aligned. Current UPMEM platforms work as accelerators, so transfers from CPU to UPMEM DIMMs (and vice versa) are needed. Additionally, DPUs lack direct communication between them. Consequently, communication between DPUs (even if they belong to the same chip) must be carefully orchestrated through the host CPU (❹/ ❺). This fact greatly impacts performance in applications where communication is frequent. While UPMEM represents current PIM architectures, PIM-capable DIMMs are expected to fully replace traditional DIMMs in future computing systems and completely remove the need for CPU-DPU data transfers.

**Key Opportunity:** UPMEM PIM platform is a suitable candidate to accelerate applications that (1) are memory-intensive, (2) require performing lightweight integer operations (i.e., addition, subtraction), (3) do not require inter-task communication. Pairwise sequence alignment algorithms met all these characteristics, including BiWFA.

Our goals are the following.

- Design and implement a PIM-enabled version of BiWFA optimized for the UPMEM PIM platform, incorporating PIM-aware optimizations to maximize performance.
- Evaluate the performance of our implementation with other state-of-the-art implementations.
- Discuss the advantages and disadvantages of the UPMEM PIM platform.

## 3 Method

### 3.1 BIMSA Overview

Sequence alignment applications usually involve comparing millions of sequence pairs. Each of these alignments can be computed independently, using their own working memory space. Therefore, BIMSA follows a coarse-grained parallelization scheme, assigning one or more sequence pairs to each DPU thread. This parallelization scheme is the best fit when targeting the UPMEM platform, as it removes the need for thread synchronization or data sharing across compute units.

We discard a collaborative parallelization scheme targeting the UPMEM platform since (1) thread synchronization and communication operations are very costly (thousands of cycles) (Gómez-Luna *et al*., 2021; Gómez-Luna *et al*., 2022; Giannoula *et al*., 2022), (2) communication between compute units has to be done through the host CPU, and(3) BiWFA’s parallelism naturally increases progressively during the alignment of a pair of sequences.

Adapting BiWFA to the UPMEM programming paradigm poses specific challenges, primarily due to BiWFA’s recursive nature and the limited stack size of UPMEM DPUs. To tackle these challenges, we opt for unrolling BiWFA’s recursion completely by (1) segmenting sequences using breakpoints, (2) generating additional BiWFA sub-alignments through iteration until a threshold is reached, and (3) generating WFA sub-alignments to finalize the alignment process.

This approach is possible since we leverage two key properties of BiWFA. First, the breakpoint iteration identifies a coordinate along the optimal path where sequences can be effectively split, separating them into two sections with balanced error. Second, each breakpoint provides the score for the iteration alignment.

Using these insights, we employ the BiWFA algorithm to iterate through sequences, breaking them down into sub-problems until the score of the sub-problem aligns with our allocated WRAM. These sub-problems, known as *base cases* in our proposal, employ the regular WFA algorithm whose memory consumption is low (i.e., dictated by the sub-problem’s remaining error rate *O*(*s*^2^)). Thus, we can precisely determine the memory required for a regular WFA within a sub-problem. Once we reach a suitable error score, we execute the WFA, producing a portion of the optimal alignment. This way, partial alignments are stored consecutively in MRAM, building progressively the complete optimal alignment.

### 3.2 PIM-Aware Optimizations

BIMSA implements nine PIM-Aware optimizations that seek to exploit the computing resources of the UPMEM platform. These optimizations illustrate effective strategies to port and accelerate memory-bound applications to the UPMEM platform.

#### Irregular Workload Balancing

Real sequencing datasets can contain sequences of different lengths and error-rate distributions. For instance, Illumina-generated datasets often have a fixed sequence length (e.g., 76 to 250 bps) and usually a low error rate (<1%)(Alser *et al*., 2022). In contrast, long-read sequencing datasets, such as `PacBio.CSS` and `Nanopore`, can contain sequence lengths ranging from hundreds of bases to thousands of bases and a larger error rate (i.e., up to 10%)(Alser *et al*., 2022).

The variety in sequence lengths and error rates rarely impacts CPU implementations regarding load balancing since the number of processing units is usually low. However, it poses two main challenges for PIM implementations where there is no dynamic memory allocation, and the number of processing units is high (e.g., 2556 in UPMEM). First, MRAM data structure allocation depends on the maximum read length in the input file for BIMSA. In cases such as `PacBio.CSS`, where most sequences fall within the 100-10 Kbps range, applications allocate space assuming the maximum sequence length. Second, restrictions on the granularity of parallelism limit the application’s performance to the slowest thread on the DPU. This scenario becomes particularly problematic when a long sequence requires more alignment time than the average, leading to an overall performance degradation.

We tackle the load balancing problem in UPMEM with three mechanisms. First, we input a maximum nominal score limit for BiWFA wavefronts, which mitigates the MRAM allocation and consumption issues and also interrupts the execution when reached. This approach involves discarding a marginal number of sequence alignments and flagging them for the CPU to complete their alignment (recovery) once the PIM kernel completes its task. Second, we assign the alignments dynamically by performing minimal synchronization based on a global alignment ID variable. Third, we execute the alignments in batches, allowing the CPU to start recovering the alignments of one batch while the PIM kernel starts computing the next batch, overlapping computations.

#### WFA Kernel Fusion

The WFA algorithm performs two core operations: compute and extend. The CPU implementation executes these two operations separately to exploit SIMD vectorization at the expense of increasing loads and stores to memory. We propose merging them into a single operation to minimize the number of MRAM access needed. To achieve this, we program manual WRAM-MRAM transfers, detecting when the last wavefront element of each WRAM block has been computed. Then, we switch to the extend operation over the WRAM block before writing it back to MRAM. In the original algorithm, computing a wavefront involves loading 3 wavefronts and writing 2. In contrast, our compute+extend fused operation reads 2 wavefronts and writes 1, effectively reducing the total MRAM accesses from 5 to 3 per wavefront element computed (i.e., reducing 40% the MRAM accesses).

#### Storing Wavefronts in MRAM

We allocate WRAM space to store the wavefronts since they are one of the most accessed data structures and we need to access them as fast as possible. However, accessing MRAM becomes necessary if the wavefronts require more than the allocated space in WRAM. Exploiting fast WRAM memory allows our implementation to compute alignments for sequences up to 100 Kbps without reducing the thread count, unlike AIM-WFA. BIMSA encounters size limitations in the MRAM memory only when handling sequences and data structures that surpass 64 MB on each DPU (i.e., MRAM’s maximum capacity). In those cases, accessing MRAM occasionally reduces performance, especially with sequences longer than 100 Kbps.

#### BiWFA Base Cases Using WRAM

To compute the alignment breakpoints, a minimum of 4 wavefronts is required (i.e., 2 for the forward and 2 for the backward alignment)(Marco-Sola *et al*., 2023). As these data structures are unnecessary for the base cases, we repurpose the allocated WRAM initially designated for these wavefronts to perform the classic WFA alignment when the alignment problem becomes sufficiently small (base cases). Hence, the base case memory requirements are calculated to fit into our predefined WRAM sizes, eliminating the need for MRAM reads and writes during their computation. This involves storing all WFA wavefronts within the space allocated for the other 4 wavefronts. As a result, the suitable error score for a base case is 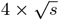, ensuring that a base case can always be entirely computed in WRAM, regardless of the size and error rate of the initial sequence.

#### Base-Case Output Reordering

During base case computation, the alignment output may have variable byte sizes that do not necessarily align with eight-byte boundaries. The UPMEM programming paradigm requires eight-byte aligned accesses to MRAM. To solve this problem, we create auxiliary WRAM and MRAM data structures that store the basecase outputs and write one output after the previous one, already writing them in the right order and storing them consecutively in the MRAM data structure. This way, when all the base cases for a sequence are computed, the complete alignment output is ready to be written back to the CPU without any concatenation operation.

#### MRAM Reverse Sequence Reading

Early versions of BIMSA involved reversing the sequences on the CPU and then transferring the duplicated and reversed sequences to the DPUs. While this simplifies memory reads on the DPU side, it comes at the cost of doubling the sequence’s MRAM usage. To tackle this problem, we implement a reverse sequence reading mechanism. This approach mitigates the risk of hitting the MRAM limit when aligning large sequences and minimizes transfer times by half.

#### Memory Align Transfer Functions

Reading a sequence in reverse retrieves WRAM blocks from the end to the start and loads data from MRAM in the opposite direction, moving from the last memory position to the initial one. However, while UPMEM offers functions to align regular reads to MRAM, these functions are not applicable to reverse reading. Consequently, we develop aligned reading functions for both forward and reverse scenarios, ensuring adaptability to different data types for the wavefront beyond 32-bit signed integers. Moreover, we adjust the entire MRAM distribution to accommodate a block’s size, ensuring compatibility in cases where reverse reading may point to an MRAM address beyond the physical limits.

#### Custom Size Transfers

BIMSA’s implementation involves many different data structures that are transferred from MRAM to WRAM with different element sizes and cadences. UPMEM allows manually allocating the WRAM space for each data structure and the transfer sizes from MRAM to WRAM. To find the best-performing configuration, we identified the cadence and size of these transfers on typical executions of BIMSA and defined the transfer sizes accordingly. The most relevant transfers are wavefront transfers and sequence transfers. However, sequence transfers read sparse data, hindering BIMSA performance when doing large reads. For this reason, we limit sequence transfers to 16 characters. Since we can not fully utilize the transfer size with sequences, we use the bigger transfer sizes on the CIGAR transfers, which occur sequentially and fully use all the memory bandwidth.

#### Adaptive Wavefront Transfer

The size of the wavefront data structure increases from the initial iteration until a breakpoint is found. Hence, initial wavefront iterations require less data movement than the last ones as the wavefronts increase in size on each algorithm iteration. We optimize UPMEM’s performance by specifying the size of data transfers from MRAM to WRAM for each iteration of the BiWFA algorithm. Initially, we set the transfer size at 8 bytes and increment it by powers of two whenever the wavefront size approaches the transfer size. We stop incrementing when the transfer size matches the size of the WRAM wavefront data structure or we reach the UPMEM transfer limit (i.e., 2048 bytes). This optimization is particularly useful when the optimal alignment score is low and most of the execution time is spent computing wavefronts smaller than the maximum transfer size, achieving a speedup of up to 1.5*×*.

## 4 Results

### 4.1 Experimental Setup

We perform the experimental evaluation on the UPMEM node described in Section 2.2 (2 Intel Xeon Silver 4215 CPUs with 8 cores each) equipped with PIM-enabled DIMM modules with 2556 operative DPUs running at 350 MHz (UPMEM-v1A). Also, we provide a projection on the performance results for the next-generation chips (UPMEM-v1B). UPMEM-v1B provides 3568 DPUs and 400Mhz per compute unit. Regarding the GPU experiments, we use an NVIDIA GeForce 3080 with 10 GB of memory. We report the average execution time of 5 runs.

We generate 8 simulated datasets for the experimental evaluation, following the methodology of WFA2lib (Marco-Sola *et al*., 2023). Each simulated dataset contains sequences of a fixed length (i.e.,5 M pairs of 150, 5 M pairs of 1 K, 2 M pairs of 10 K, and 12 K pairs of 100 Kbps), error rate (i.e., 5% and 10%). Additionally, we present an evaluation using real sequencing datasets, described in Section 2.2, which contain sequences of different lengths and error rates, representing heterogeneous and irregular workloads.

We evaluate and compare the performance of BIMSA by selecting the already presented CPU application WFA2lib, the state-of-the-art GPU implementation of the WFA (Aguado-Puig *et al*., 2023) and the PIM library AIM (Diab *et al*., 2023), which implements a gap-affine optimized Needleman-Wunsch (NW) algorithm, the classical WFA algorithm for PIM and an adaptive heuristic for WFA. For simulated datasets, we evaluate BIMSA in the current UPMEM chips (UPMEM-v1A) and also provide a performance projection of the next-generation chips (UPMEM-v1B) based on the scalability results. For PIM executions, we report kernel execution time, assuming data is loaded in the device’s memory.

The CPU implementation is executed with the maximum number of hardware threads (16 threads) exploiting coarse grain parallelism. All the PIM implementations are executed with 2500 DPUs. BIMSA is executed using 12 tasklets. The number of tasklets used by AIM is set by their execution script and is lowered if the data cannot fit the WRAM.

### 4.2 BIMSA Characterization Using Simulated Datasets

Figure 3 illustrates the performance, in alignments per second, achieved by the state-of-the-art applications and BIMSA across balanced workload datasets. The numbers on top of each bar represent the speedup needed to match the fastest application.

**Fig. 3:**
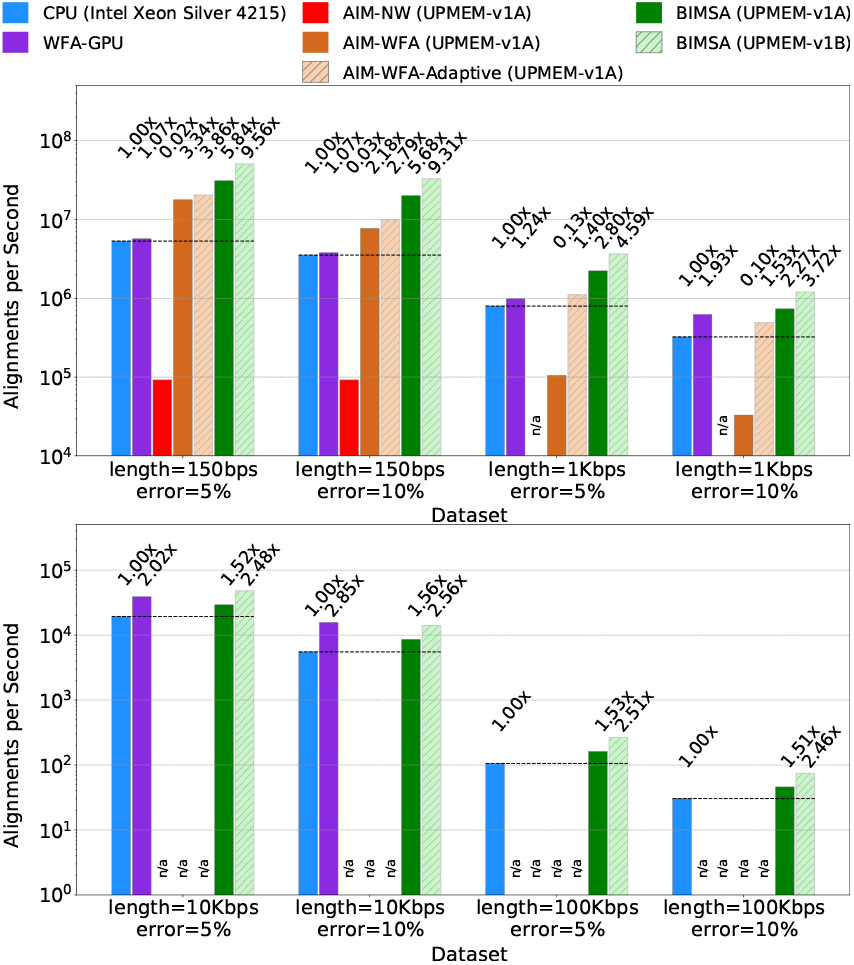
Alignments per second achieved by WFA2lib (CPU), WFA-GPU (GPU), AIM (PIM), and BIMSA (PIM) when aligning simulated datasets. Failed or unsupported experiments are marked as n/a.

First, we observe that AIM-NW exhibits lower performance than the rest of the evaluated applications and fails to align long sequence datasets. This is expected since AIM-NW is based on a traditional DP algorithm that requires a large memory footprint.

Second, we observe that BIMSA outperforms the PIM state-of-the-art (AIM-WFA) across all datasets, achieving speedups up to 22.24*×* (11.95*×* on average). Additionally, while BIMSA handles sequences of 10 Kbps and longer, AIM-WFA fails due to insufficient memory allocation space. Third, we observe that AIM-WFA-Adaptive obtains up to 14.9*×* speedup (7*×* on average) compared to the regular AIM-WFA version while producing 100% accurate results for all datasets (using the default heuristic parameters) as shown in the Supplementary Section S1. However, it is still outperformed by BIMSA up to 2*×* (1.75*×* on average) and fails to handle sequences of 10 Kbps or longer.

Fourth, the GPU-WFA implementation is outperformed by AIM-WFA and BIMSA on the 150 sequence length datasets. However, it outperforms AIM-WFA on the 1 Kbps sequences and can handle larger sequences than AIM-WFA (up to 10 K). Additionally, in the 10 Kbps sequences, WFA-GPU is able to outperform BIMSA by up to 1.8*×*. Nevertheless, BIMSA outperforms WFA-GPU in all the other datasets up to 5.4*×* (2.5*×* on average) and can handle larger sequences (100 Kbps). This is due to WFA-GPU being programmed with 16 bit wavefront elements, which limits the sequence length to 30 Kbps.

Fifth, comparing BIMSA to the state-of-the-art CPU performance, we observe that BIMSA outperforms CPU implementations by up to 5.84*×* (2.83*×* on average). However, we note that as the sequence length and error percentage increase, BIMSA’s performance degrades, achieving 1.51*×* speedup in the worst-case scenario. This is partly because the highly sequential WFA extend operations become a major bottleneck. Nevertheless, by projecting these results using the scalability results from Figure 4, we observe that BIMSA outperforms CPU implementations by up to 9.56*×* (4.7*×* in average) and at least 2.5*×* in the worst case.

**Fig. 4:**
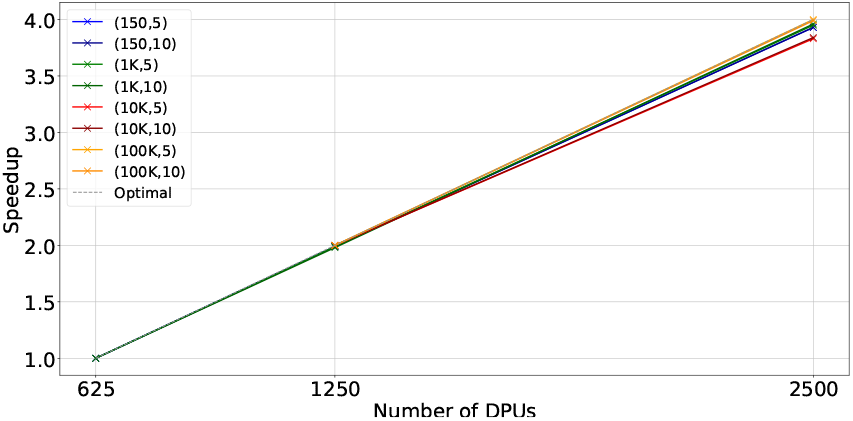
Scalability with the number of DPUs for BIMSA with balanced workload datasets. The datasets are indicated by (length, error %).

Table 2 shows the execution times for the datasets and applications presented in Figure 3. For the PIM-based experiments, results that include PIM transfer times are marked with (+T). We observe that the transfer times reduce the performance of PIM applications in short sequences but are negligible in longer sequences. However, we evaluate all cases without considering transfer times due to the forecasting of upcoming PIM systems that do not require such transfers.

**Table 2.** Execution time in seconds of WFA2lib (CPU), WFA-GPU (GPU), AIM (PIM), and BIMSA (PIM). For the PIM-based experiments, results that include PIM transfer times are marked with (+T). Failed or unsupported experiments are marked as n/a.

|  | l=150 |  | l=1K |  | l=10K |  | l=100K |  |
| --- | --- | --- | --- | --- | --- | --- | --- | --- |
|  | e=5% | e=10% | e=5% | e=10% | e=5% | e=10% | e=5% | e=10% |
| WFA2lib | 0.9 | 1.4 | 6.3 | 15.5 | 103.6 | 363.2 | 568.2 | 1970.4 |
| WFA-GPU | 0.9 | 1.3 | 5.1 | 8.0 | 51.4 | 127.6 | n/a | n/a |
| BIMSA | 0.2 | 0.3 | 2.3 | 7.0 | 69.9 | 238.4 | 371.2 | 1312.4 |
| BIMSA (+T) | 2.5 | 2.6 | 6.4 | 11.0 | 79.7 | 243.8 | 378.6 | 1319.6 |
| AIM-WFA | 0.3 | 0.6 | 47.5 | 151.6 | n/a | n/a | n/a | n/a |
| AIM-WFA (+T) | 1.7 | 2.0 | 50.2 | 154.3 | n/a | n/a | n/a | n/a |
| AIM-WFA-Adapt | 0.2 | 0.5 | 4.5 | 10.1 | n/a | n/a | n/a | n/a |
| AIM-WFA-Adapt (+T) | 1.7 | 2.0 | 7.8 | 13.6 | n/a | n/a | n/a | n/a |
| AIM-NW | 54.3 | 54.4 | n/a | n/a | n/a | n/a | n/a | n/a |
| AIM-NW (+T) | 57.2 | 57.3 | n/a | n/a | n/a | n/a | n/a | n/a |

**Key Result I:** BIMSA outperforms the state-of-the-art PIM implementations by up to 22.24*×* and state-of-the-art CPU by up to 5.84*×* when dealing with simulated datasets. Furthermore, the projection on upcoming UPMEM-v1B systems achieves speedups up to 9.56*×*.

In Figure 4, we examine the scalability of the same datasets as in Figure 3 across different numbers of DPUs. A dashed gray line denotes ideal scalability. We observe that BIMSA’s scalability with the number of DPUs is nearly linear, while CPU implementations do not scale well when increasing the number of cores (as demonstrated in Section 2.2, Figure 1). It is expected that next-generation UPMEM systems will double the number of DPUs. Thanks to the linear scalability of BIMSA, this will translate into significant performance improvements.

**Key Result II:** BIMSA exhibits linear scalability with the number of compute units.

In Figure 5, we analyze the scalability of BIMSA and the same datasets as in Figure 4 by varying the number of threads and fixing the number of DPUs to 2500. We make three observations from it.

**Fig. 5:**
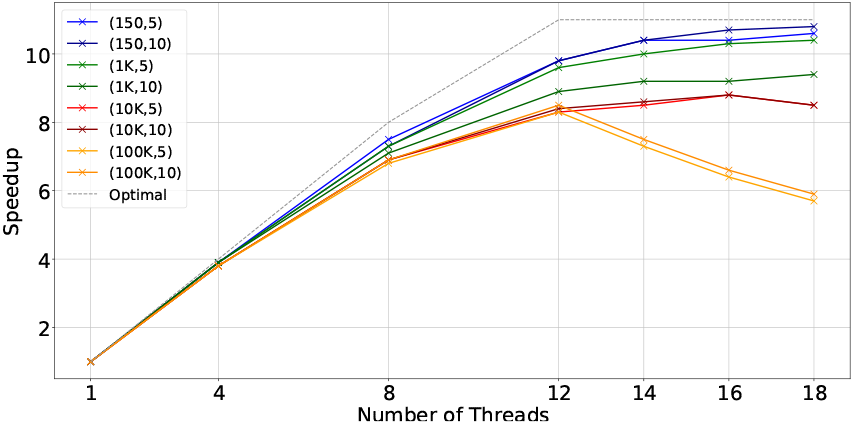
Scalability with the number of threads for BIMSA with balanced workload datasets. The datasets are indicated by (length, error %).

First, we notice that BIMSA struggles to scale between 12 and the maximum number of threads. This is an expected behavior and matches a limitation of the UPMEM architecture. In this sense, prior work (Gómez-Luna *et al*., 2022) has demonstrated that PIM applications that fully utilize resources typically scale only up to 11 threads. This limit arises because each thread of the DPU processor can dispatch only one instruction every 11 cycles due to the pipeline depth.

Second, we find that as the sequence size increases, BIMSA’s scalability deteriorates with a lower number of threads. This is attributed to the increase in MRAM accesses for long sequence lengths. Since MRAM can only be accessed by one thread at a time, threads encounter a bottleneck in MRAM access, particularly noticeable with larger sequence sizes.

Third, we can see how datasets with 100 Kbps per sequence lose scalability and performance regarding the 12 thread execution. This is because the number of pairs is insufficient to provide even work for all the threads, which shows how unbalanced work hinders UPMEM’s performance.

### 4.3 BIMSA Characterization Using Real Datasets

Table 3 illustrates the execution time in seconds achieved by the state-of-the-art applications across different unbalanced workload datasets. In this analysis, BIMSA consists of two versions: first, an implementation that executes all the alignments in the DPUs (*BIMSA*), and second, an implementation that executes alignments in the DPUs up to a maximum alignment-score threshold and then recovers the alignments with higher alignment-score in the CPU (*BIMSA-Hybrid*). In particular, the recovery version uses one CPU thread to manage DPU requests and 16 threads to execute the CPU alignments, exploiting a coarse-grain parallelism approach (i.e., one alignment per thread). To evaluate this version, we measure the time since the first alignment kernel execution until all the alignments have finished both in DPU and CPU.

**Table 3.**
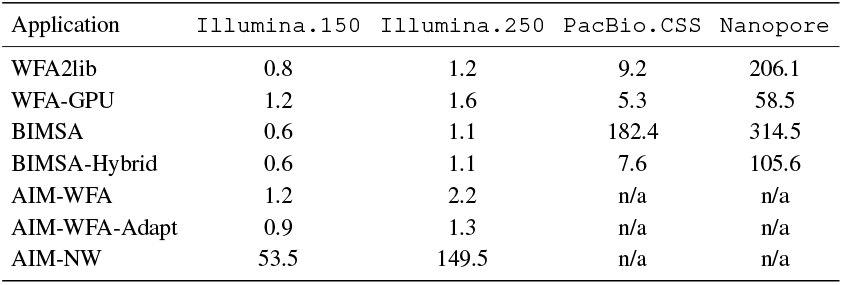
Execution time in seconds of AIM (PIM), WFA2lib (CPU), WFA-GPU, BIMSA (PIM) and BIMSA-Hybrid (PIM) when aligning unbalanced real datasets. Failed or unsupported experiments are marked as n/a.

We show that BIMSA outperforms the prior state-of-the-art PIM (AIM) across all scenarios, achieving up to 2.08*×* speedup and processing even the longest sequences datasets that the AIM library cannot. AIM-WFA-Adaptive outperforms AIM-WFA producing results 99.99% accurate, as shown in the Supplementary Section S1. However, it is still outperformed by BIMSA by up to 1.5*×*. Regarding CPU performance, we observe that BIMSA provides similar performance for the `Illumina` datasets in both versions. Looking at BIMSA when processing long-sequence datasets, we can see how we outperform other CPU implementations aligning `PacBio.CSS` and have similar performance in `Nanopore`. In particular, the CPU recovery optimization (i.e., BIMSA + CPU) helps to reduce the DPU’s execution time variability between alignments, making BIMSA able to outperform WFA2lib by 1.21*×* times in `PacBio.CSS` and 1.95*×* in `Nanopore` by aligning 7.36% and 12.75% of pairs in the CPU respectively. Note that Illumina datasets are more homogeneous than `PacBio.CSS` and `Nanopore`, resulting in the same execution for BIMSA and BIMSA-Hybrid. In contrast, we observe that WFA-GPU is able to outperform all the PIM-implementations in the `PacBio.CSS` and `Nanopore` datasets. Compared to BIMSA-Hybrid, WFA-GPU achieves speedups of 1.4*×* and 1.8*×* for the `PacBio.CSS` and `Nanopore` datasets, respectively. This shows the limitation that load unbalance represents for UPMEM applications. Regardless, BIMSA outperforms WFA-GPU on the `Illumina` datasets up to 2*×*.

**Key Result III:** BIMSA outperforms state-of-the-art PIM solutions by up to 2.08*×* and state-of-the-art CPU implementations by up to 1.95*×* when dealing with irregular sequencing datasets thanks to its balancing and recovery method.

## 5 Discussion

In this work, we introduce BIMSA, a PIM-based implementation of the BiWFA sequence alignment algorithm. BIMSA leverages the real-world general-purpose PIM architecture developed by UPMEM and designed to address data-movement inefficiencies that limit the scalability of sequence alignment implementations.

### Key outcomes of BIMSA

Our evaluation shows that, when aligning synthetic datasets, BIMSA achieves speedups up to 22.24*×* (11.95*×* on average) compared to state-of-the-art PIM-enabled implementations of sequence alignment algorithms while also processing sequence lengths beyond the limitations of the PIM-enabled state-of-the-art. BIMSA achieves speedups up to 5.84*×* (2.83*×* on average) compared to the BiWFA CPU implementation. BIMSA achieves speedups up to 9.56*×* (4.7*×* on average) in the UPMEM-v1B chips projection. Similarly, when aligning real datasets, BIMSA achieves speedups up to 2.08*×* compared to the PIM-based AIM library and 1.95*×* compared to BiWFA CPU. These results demonstrate that BIMSA performs better than prior state-of-the-art sequence alignment implementations. Most importantly, we observe ideal scalability with the number of compute units that gives BIMSA the potential of fully utilizing upcoming PIM systems.

### Limitations of BIMSA

We observe two key limitations of BIMSA. First, there is a performance degradation when dealing with high error sequences (starting at 10 K bases 5% error). This is explained by the fact that the size of the wavefronts is proportional to the error rate. Consequently, when the wavefront size exceeds the allocated WRAM space, the number of WRAM-MRAM wavefront transfers increases and the execution performance is reduced. Second, processing heterogeneous datasets (i.e., sequence pairs of different lengths and error-rates) generates load unbalance between compute units. This requires careful workload distribution across a large number of available compute units, which are unable to perform efficient inter-DPU communication.

### Potential of PIM for Long Sequence Alignment

Since PIM systems are at an early stage of technological development, there are many opportunities for improvement in the upcoming generations. We anticipate four key features that will further help accelerate core bioinformatics algorithms.

- *Direct Communication Between Host and DPUs*. This feature would allow the system to remove the need for transfers and also would enable more efficient CPU-DPU collaboration. In particular, BIMSA can recover large alignments in the CPU. However, it needs to (1) wait for other alignments to finish and (2) start the alignment from scratch on the CPU. Direct CPU-DPU communication would allow BIMSA to resume the alignment once it is interrupted in the DPUs, while it keeps working on new alignments.
- *Efficient DPU intercommunication*. This feature would enable workload sharing across DPUs that have finished their alignments before other DPUs, alleviating the work unbalance, a major UPMEM limitation.
- *Support for Vector Instructions*. This feature would help accelerate compute-bound sections of bioinformatics applications. For instance, the compute step of BIMSA would benefit from this feature by simultaneously computing multiple wavefront elements using a single SIMD instruction.
- *Support for DPX-like instructions*. In the same spirit as the DPX instruction set implemented on the latest GPU models (Luo *et al*., 2024), we believe these instructions would provide significant acceleration to applications like BIMSA running in UPMEM. For instance, one of BIMSA’s core operations involves calculating the maximum between three values. In the current UPMEM architecture, this operation requires 10 instructions; while using a DPX-like instruction, the number of required instructions would drop to only 6.

Overall, this work demonstrates that PIM technology is a suitable solution for accelerating sequence alignment, even considering its early stages of development. Considering the ongoing development of PIM technology and the numerous possibilities for this emerging architecture, we argue that PIM-based solutions are a promising choice for accelerating sequence alignment and other memory-bound bioinformatics applications. Notwithstanding, PIM-based solutions are not yet ready for production environments. This is mainly due to the lack of a mature ecosystem of PIM-enabled bioinformatics applications. Also, developing PIM-accelerated applications still requires significant time and effort until the PIM development toolchain improves. Recent PIM-based solutions, such as BIMSA, require further integration and testing into full-fledged bioinformatics data analysis pipelines before the bioinformatics community can take full advantage of this technology. Regardless, we are confident in the potential of PIM-based solutions to effectively accelerate future bioinformatics applications.

## References

(2022). 184QPS/W 64Mb/mm 2 3D Logic-to-DRAM Hybrid Bonding With Process-Near-Memory Engine for Recommendation System, author=Niu, Dimin and Li, Shuangchen and Wang, Yuhao and Han, Wei and Zhang, Zhe and Guan, Yijin and Guan, Tianchan and Sun, Fei and Xue, Fei and Duan, Lide and others. In 2022 IEEE International Solid-State Circuits Conference (ISSCC), volume 65, pages 1–3. IEEE.

Aguado-Puig, Q., Marco-Sola, S., Moure, J. C., Castells-Rufas, D., Alvarez, L., Espinosa, A., and Moreto, M. (2022). Accelerating Edit-Distance Sequence Alignment on GPU Using the Wavefront Algorithm. IEEE access, 10, 63782– 63796.

Aguado-Puig, Q., Doblas, M., Matzoros, C., Espinosa, A., Moure, J. C., Marco-Sola, S., and Moreto, M. (2023). Wfa-gpu: gap-affine pairwise read-alignment using gpus. Bioinformatics, 39(12), btad701.

Alser, M., Rotman, J., Deshpande, D., Taraszka, K., Shi, H., Baykal, P. I., Yang, H. T., Xue, V., Knyazev, S., Singer, B. D., et al. (2021). Technology dictates algorithms: recent developments in read alignment. Genome biology, 22(1), 249.

Alser, M., Lindegger, J., Firtina, C., Almadhoun, N., Mao, H., Singh, G., Gomez-Luna, J., and Mutlu, O. (2022). From Molecules to Genomic Variations: Accelerating Genome Analysis Via Intelligent Algorithms and Architectures. Computational and Structural Biotechnology Journal.

Balhaf, K., Shehab, M. A., Wala’a, T., Al-Ayyoub, M., Al-Saleh, M., and Jararweh, Y. (2016). Using GPUs to Speed-Up Levenshtein Edit Distance Computation. In 2016 7th International Conference on Information and Communication Systems (ICICS), pages 80–84. IEEE.

Churko, J. M., Mantalas, G. L., Snyder, M. P., and Wu, J. C. (2013). Overview of High Throughput Sequencing Technologies to Elucidate Molecular Pathways in Cardiovascular Diseases. Circulation research, 112(12), 1613–1623.

Daily, J. (2016). Parasail: SIMD C Library for Global, Semi-Global, and Local Pairwise Sequence Alignments. BMC bioinformatics, 17(1), 1–11.

Diab, S., Nassereldine, A., Alser, M., Gómez Luna, J., Mutlu, O., and El Hajj, I. (2023). A Framework for High-Throughput Sequence Alignment Using Real Processing-in-Memory Systems. Bioinformatics, 39(5), btad155.

Gerometta, G., Zeni, A., and Santambrogio, M. D. (2023). TSUNAMI: A GPU Implementation of the WFA Algorithm. In 2023 32nd International Conference on Parallel Architectures and Compilation Techniques (PACT), pages 150–161. IEEE.

Ghose, S., Boroumand, A., Kim, J. S., Gómez-Luna, J., and Mutlu, O. (2019). Processing-in-memory: A Workload-Driven Perspective. IBM Journal of Research and Development, 63(6), 3–1.

Giannoula, C., Fernandez, I., Luna, J. G., Koziris, N., Goumas, G., and Mutlu, O. (2022). Sparsep: Towards efficient sparse matrix vector multiplication on real processing-in-memory architectures. Proceedings of the ACM on Measurement and Analysis of Computing Systems, 6(1), 1–49.

Gómez-Luna, J., El Hajj, I., Fernandez, I., Giannoula, C., Oliveira, G. F., and Mutlu, O. (2021). Benchmarking Memory-Centric Computing Systems: Analysis of real Processing-In-Memory Hardware. In 2021 12th International Green and Sustainable Computing Conference (IGSC), pages 1–7. IEEE.

Gotoh, O. (1982). An Improved Algorithm for Matching Biological Sequences. Journal of molecular biology, 162(3), 705–708.

Gómez-Luna, J., Hajj, I. E., Fernandez, I., Giannoula, C., Oliveira, G. F., and Mutlu, O. (2022). Benchmarking a New Paradigm: Experimental Analysis and Characterization of a Real Processing-in-Memory System. IEEE Access, 10, 52565–52608.

Haghi, A., Marco-Sola, S., Alvarez, L., Diamantopoulos, D., Hagleitner, C., and Moreto, M. (2021a). An FPGA Accelerator of the Wavefront Algorithm for Genomics Pairwise Alignment. In 2021 31st International Conference on Field-Programmable Logic and Applications (FPL), pages 151–159. IEEE.

Haghi, A., Marco-Sola, S., Alvarez, L., Diamantopoulos, D., Hagleitner, C., and Moreto, M. (2021b). An fpga accelerator of the wavefront algorithm for genomics pairwise alignment. In 202131st International Conferenceon Field-Programmable Logic and Applications (FPL), pages 151–159.

Haghi, A., Alvarez, L., Front, J., De Haro Ruiz, J.M., Figueras, R., Doblas, M., Marco-Sola, S., and Moreto, M. (2023). WFAsic: A High-Performance ASIC Accelerator for DNA Sequence Alignment on a RISC-V SoC. In Proceedings of the 52nd International Conference on Parallel Processing, pages 392–401.

Hajinazar, N., Oliveira, G. F., Gregorio, S., Ferreira, J., Ghiasi, N. M., Patel, M., Alser, M., Ghose, S., Luna, J. G., and Mutlu, O. (2021). SIMDRAM: An End-to-End Framework for bit-serial SIMD Computing in DRAM. arXiv preprint arXiv:2105.12839.

Intel (2019). Intel Xeon Silver 4215. https://www.intel.com/content/www/us/en/products/sku/193389/intel-xeon-silver-4215-processor-11m-cache-2-50-ghz/specifications.html.

Kautz, W. H. (1969). Cellular logic-in-memory arrays. IEEE Transactions on Computers, 100(8), 719–727.

Ke, L., Zhang, X., So, J., Lee, J.-G., Kang, S.-H., Lee, S., Han, S., Cho, Y., Kim, J. H., Kwon, Y., et al. (2021). Near-Memory Processing in Action: Accelerating Personalized Recommendation With Axdimm. IEEE Micro, 42(1), 116–127.

Kwon, Y.-C., Lee, S. H., Lee, J., Kwon, S.-H., Ryu, J. M., Son, J.-P., Seongil, O., Yu, H.-S., Lee, H., Kim, S. Y., et al. (2021). 25.4 a 20nm 6Gb Function-in-Memory DRAM, Based on HBM2 With a 1.2 Tflops Programmable Computing Unit Using Bank-Level Parallelism, for Machine Learning Applications. In 2021 IEEE International Solid-State Circuits Conference (ISSCC), volume 64, pages 350–352. IEEE.

Lee, S., Kang, S.-h., Lee, J., Kim, H., Lee, E., Seo, S., Yoon, H., Lee, S., Lim, K., Shin, H., et al. (2021). Hardware Architecture and Software Stack for PIM Based on Commercial DRAM Technology: Industrial Product. In 2021 ACM/IEEE 48th Annual International Symposium on Computer Architecture (ISCA), pages 43–56. IEEE.

Lee, S., Kim, K., Oh, S., Park, J., Hong, G., Ka, D., Hwang, K., Park, J., Kang, K., Kim, J., et al. (2022). A 1ynm 1.25 v 8Gb, 16Gb/s/pin GDDR6-based Accelerator-in-Memory Supporting 1Tflops Mac Operation and Various Activation Functions for Deep-Learning Applications. In 2022 IEEE International Solid-State Circuits Conference (ISSCC), volume 65, pages 1–3., pages IEEE.

Luo, W., Fan, R., Li, Z., Du, D., Wang, Q., and Chu, X. (2024). Benchmarking and dissecting the nvidia hopper gpu architecture. arXiv preprint arXiv:2402.13499.

Marco-Sola, S., Moure, J. C., Moreto, M., and Espinosa, A. (2021). Fast Gap-Affine Pairwise Alignment Using the Wavefront Algorithm. Bioinformatics, 37(4), 456–463.

Marco-Sola, S., Eizenga, J. M., Guarracino, A., Paten, B., Garrison, E., and Moreto, M. (2023). Optimal Gap-Affine Alignment in O(s) Space. Bioinformatics, 39(2), btad074.

Mutlu, O., Ghose, S., Gómez-Luna, J., and Ausavarungnirun, R. (2019a). Enabling Practical Processing in and Mear Memory for Data-Intensive Computing. In Proceedings of the 56th Annual Design Automation Conference 2019, pages 1–4.

Mutlu, O., Ghose, S., Gómez-Luna, J., and Ausavarungnirun, R. (2019b). Processing Data Where it Makes Sense: Enabling In-Memory Computation. Microprocessors and Microsystems, 67, 28–41.

Mutlu, O., Ghose, S., Gómez-Luna, J., and Ausavarungnirun, R. (2022). A Modern Primer on Processing in Memory. In Emerging Computing: From Devices to Systems: Looking Beyond Moore and Von Neumann, pages 171–243. Springer.

Needleman, S. B. and Wunsch, C. D. (1970). A General Method Applicable to the Search for Similarities in the Amino Acid Sequence of two Proteins. Journal of molecular biology, 48(3), 443–453.

NIST (2023). Giab Data Indexes.https://github.com/genome-in-a-bottle/giab_data_indexes.

Reuter, J. A., Spacek, D. V., and Snyder, M. P. (2015). High-Throughput Sequencing Technologies. Molecular cell, 58(4), 586–597.

Roy, S., Coldren, C., Karunamurthy, A., Kip, N. S., Klee, E. W., Lincoln, S. E., Leon, A., Pullambhatla, M., Temple-Smolkin, R. L., Voelkerding, K. V., et al. (2018). Standards and Guidelines for Validating Next-Generation Sequencing Bioinformatics Pipelines: A Joint Recommendation of the Association for Molecular Pathology and the College of American Pathologists. The Journal of Molecular Diagnostics, 20(1), 4–27.

Schloss, J. A. (2008). How to Get Genomes at one Ten-Thousandth the Cost. Nature biotechnology, 26(10), 1113–1115.

Stone, H. S. (1970). A Logic-in-Memory Computer. IEEE Transactions on Computers, 100(1), 73–78.

UPMEM (2024). UPMEM Website. https://www.upmem.com.

Vasimuddin, M., Misra, S., Li, H., and Aluru, S. (2019). Efficient Architecture-Aware Acceleration of BWA-MEM for Multicore Systems. In 2019 IEEE international parallel and distributed processing symposium (IPDPS), pages 314–324. IEEE.

Walia, S., Ye, C., Bera, A., Lodhavia, D., and Turakhia, Y. (2024). TALCO: Tiling Genome Sequence Alignment Using Convergence of Traceback Pointers. In 2024 IEEE International Symposium on High-Performance Computer Architecture (HPCA), pages 91–107. IEEE.

Waterman, M. S., Smith, T. F., and Beyer, W. A. (1976). Some Biological Sequence Metrics. Advances in Mathematics, 20(3), 367–387.

