## Supplementary Material for "BIMSA: Accelerating Long Sequence Alignment Using Processing-In-Memory"

### S1 AIM-WFA-Adaptive Accuracy Percentage

In this section, we present the accuracy results for the WFA heuristic implementation AIM-WFA-Adaptive experiments. In this analysis, accuracy refers to the number of optimal alignments reported by the tool. Table S1 presents the AIM-WFA-Adaptive accuracy percentage for every dataset aligned. The experiments use the default parameters, which provide at least 99.99% accuracy for all evaluated datasets.

| **Dataset** | **Accuracy %** |
|---|---|
| (l=150, e=5%) | 100.00% |
| (l=150, e=10%) | 100.00% |
| (l=1000, e=5%) | 100.00% |
| (l=1000, e=10%) | 100.00% |
| (l=10000, e=5%) | Not supported |
| (l=10000, e=10%) | Not supported |
| (l=100000, e=5%) | Not supported |
| (l=100000, e=10%) | Not supported |
| Illumina.150.10M | 99.99% |
| Illumina.250.10M | 99.99% |
| PacBio.CSS.1M | Not supported |
| Nanopore.Bowden.1M | Not supported |

Table S1: Accuracy percentage of AIM-WFA-Adaptive for each dataset. Failed executions or not supported by AIM-WFA-Adaptive due to a lack of WRAM space are marked as "Not supported".

### S2 WFA and BiWFA Performance Characterization

This section presents a detailed CPU performance characterization of WFA and BiWFA. Both algorithms use the same functions to compute successive wavefront vectors until the alignment problem is solved, known as *Compute Wavefront* and *Extend Wavefront*. Wavefronts dynamically expand as the alignment progress continues until its completion.

During the first stages of the alignment, wavefront vectors are small and fit in the CPU caches, putting low pressure on the memory subsystem. As the alignment algorithm continues, the wavefront's size increases beyond the CPU Last Level Cache (LCC) capacity, requiring access to the main memory. As a result, the pressure on the memory subsystem increases. Similarly, as the number of working threads in the CPU increases, the LLC and main memory have to handle more memory requests. In turn, this puts more pressure on the memory subsystem.

To analyze this behaviour, we implement a multi-threaded benchmark that analyzes the performance of the WFA and BiWFA algorithms at any given step of the alignment process. For that, our benchmark executes repeatedly the *Compute Wavefront* and *Extend Wavefront* functions over a wavefront of a fixed size. Moreover, this benchmark can spawn multiple threads computing wavefronts on different processor cores to reproduce the algorithm's performance executed on a multicore processor. During the algorithm's execution, the benchmark collects CPU performance counters to profile the applications' performance. Each step is performed repeatedly to obtain an accurate measurement and reduce system interference.

We run this benchmark for different wavefront sizes, ranging from 1 KiB up to 4 MiB. Each thread works on a private wavefront and sequence pair, simulating the real-case scenario where each thread aligns a different sequence pair. The UPMEM platform has 16 physical cores, so we conduct experiments with 1, 4, 8, and 16 threads. We perform three sets of experiments using this benchmark.

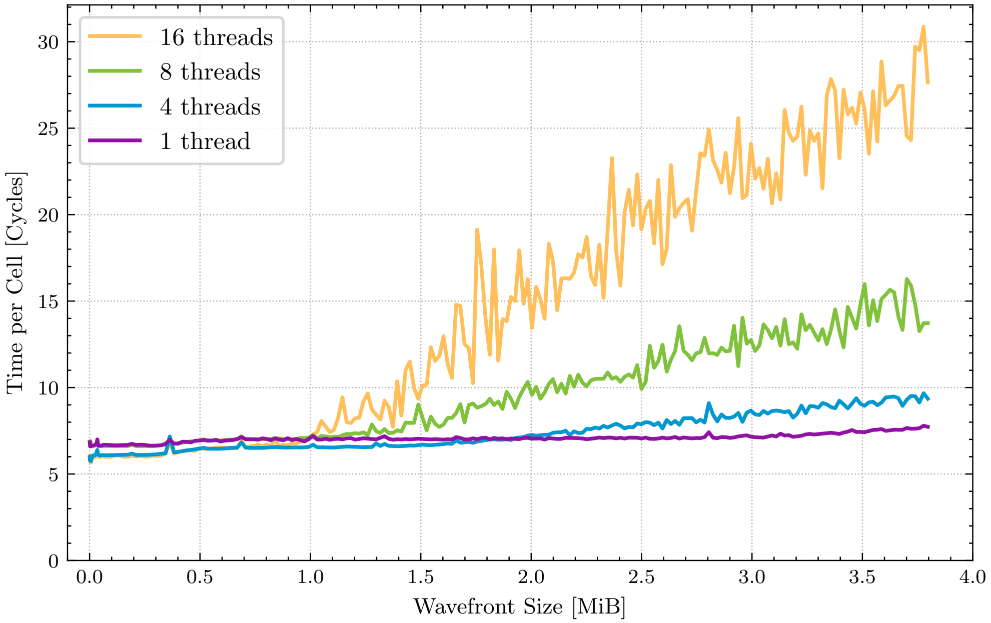


Figure S1: Execution cycles per wavefront element computed varying the wavefront size and the number of threads.

First, we analyze the average time taken to compute each wavefront element, varying the wavefront size and the number of threads. The result of this analysis is shown in Fig. S1. We observe that the time per cell grows when computing wavefronts bigger than 1 MiB. Moreover, the time required per cell increases as more threads are utilised. This indicates that the memory subsystem is under high pressure and potentially saturated due to the high volume of concurrent requests to the LCC and memory.

Second, we measure LCC references and LLC misses (i.e., the number of LCC requests that require DRAM access). This experiment evaluates whether the memory subsystem is saturated, characterizing the algorithm as memory-bound. The results are shown in Fig. S2. We observe that the number of references to the LLC increases when wavefronts become bigger than than 1 MiB and stay the same regardless of the number of threads. This is expected when wavefronts exceed

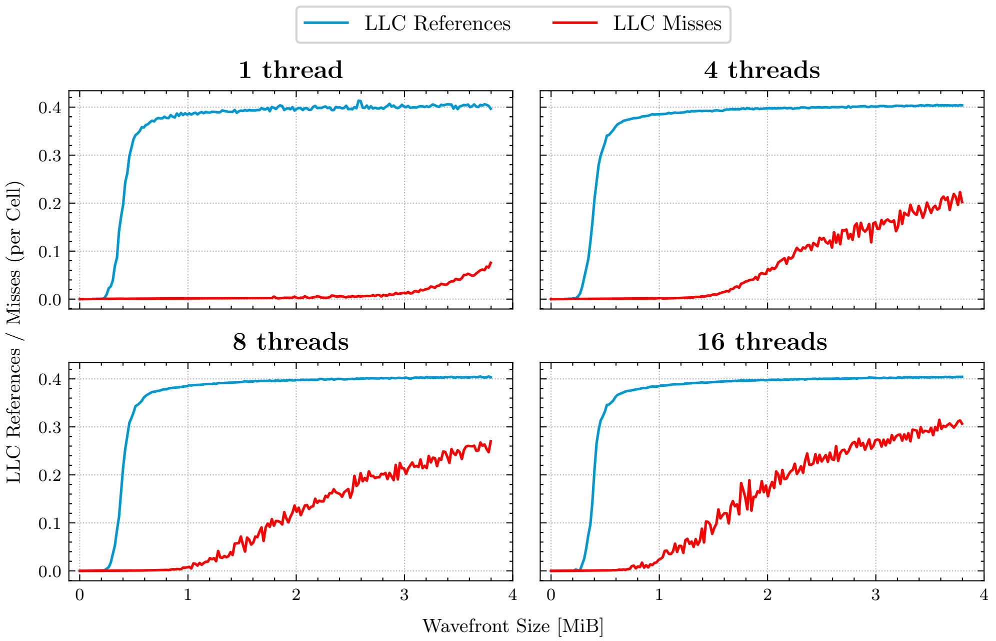


Figure S2: Average LLC references and misses per wavefront element computed varying the wavefront size and the number of threads.

L2 capacity, forcing the CPU to access the LLC for data. However, we observe differences in the LLC misses. Using a single thread, there are almost no cache misses on the LLC until very large wavefronts are processed. This indicates that most of the data is accessed from the L1 or L2 cache, which is consistent with the results from our previous experiment. In contrast, executing with more threads generates conflicts in the LLC, increasing the number of LLC misses and limiting the performance due to memory requests saturation.

Third, we measure the MPKI (LLC Misses Per Kilo Instructions) in Fig. S3. MPKI is a commonly used metric to evaluate the cache's performance when executing a specific program. It is calculated using Eq. 1. A low MPKI value indicates good memory cache usage, as cache misses are relatively low compared with the number of instructions executed. In general, MPKI can be divided into different ranges that indicate good cache usage (0-5), average cache usage (5-10), and poor cache usage (>10). We observe that the executions of 4, 8, and 16 threads fall into the average and poor performance.

$$\text{MPKI} = \frac{\text{LLC Misses}}{\text{Instructions}} \times 1000 \tag{1}$$

These three experiments demonstrate that WFA and BiWFA are predominantly *memory bound* in modern CPUs as the wavefront size and number of working threads increase. In turn, these results demonstrate that Processing-In-Memory is a promising solution to mitigate these performance bottlenecks.

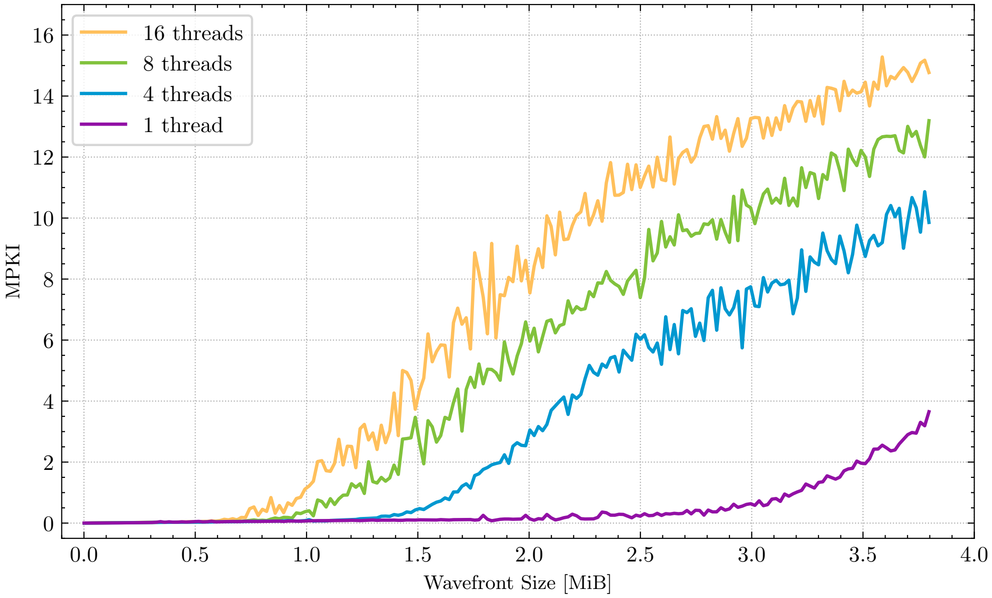


Figure S3: LLC MPKI (Last Level Cache Misses per Kilo Instructions) varying the wavefront size and the number of threads.
